# Stable flies are bonafide reservoirs of mastitis-associated bacteria

**DOI:** 10.1101/2024.02.27.582344

**Authors:** Andrew J. Sommer, Julia E. Kettner, Kerri L. Coon

## Abstract

Hematophagous *Stomoxys* (stable) fly populations in dairy barns are sustained by a constant availability of cattle hosts and manure, which serve as major reservoirs of both zoonotic and opportunistic bacterial pathogens. However, the overall composition and diversity of bacterial communities associated with *Stomoxys* flies and the ability of biting flies to acquire and transmit potentially pathogenic bacteria present in their surrounding environment remain to be investigated. Here, we present the first culture-independent examination of *Stomoxys*-associated bacterial communities through longitudinal sampling of fly and manure samples collected from two connected dairy facilities in South Central Wisconsin. High throughput 16S rRNA gene amplicon sequencing was used to characterize and compare bacterial communities present on or within flies and in manure collected from the same facility. Bacterial alpha diversity was overall higher in manure samples as compared to fly samples, with manure-associated bacterial communities being dominated by members of the Bacteroidales, Eubacteriales, and Oscillospirales. In contrast, flies harbored relatively low-complexity communities dominated by members of the Enterobacterales, Staphylococcales, and Lactobacillales. Clinically relevant bacterial strains, including *Escherichia* spp. and other taxa associated with mastitic cows housed in the same facilities, were detected in paired fly and manure samples but exhibited dramatically elevated abundances in fly samples as compared to manure samples. Viable colonies of *Escherichia*, *Klebsiella*, and *Staphylococcus* spp. were also readily isolated from fly samples, confirming that flies harbor culturable mastitis associated bacteria. This study provides definitive support for a potential role for biting flies in mediating bacterial pathogen transmission in dairy barns and other agricultural settings.

**IMPORTANCE:** Disease prevention on dairy farms has significant implications for cattle health, food security, and zoonosis. Of particular importance is the control of bovine mastitis, which can be caused by a diverse array of environmental bacterial pathogens, including *Klebsiella*, *E. coli*, *Streptococcus*, and *Staphylococcus* spp. Despite being one of the most significant and costly cattle diseases worldwide, the epidemiology of bovine mastitis is not well understood. This study provides the first culture-dependent and culture-independent evidence to support the potential for biting flies to transmit opportunistic bovine and human pathogens in agricultural settings. It also links carriage of specific bacterial taxa in flies to clinical mastitis cases in cows housed in the same facility at the time of sampling. Altogether, these results indicate that biting flies represent an important, yet understudied biosecurity threat to animal husbandry facilities.

## INTRODUCTION

Bovine mastitis is widely considered to be one of the most significant and widespread diseases facing the dairy industry, with the USDA National Animal Health Monitoring System (NAHMS) reporting cases in 99.7% of all U.S. dairy operations (USDA, 2018). The economic losses associated with mastitis cases are especially burdensome, costing the dairy industry an estimated 35 billion dollars annually (National Mastitis Council, 2016). Not only does low-quality milk need to be discarded, but dairy farmers must also pay for costs associated with lost labor, as well as the treatment and potential culling of sick cows (Halasa et al., 2007). Symptoms of bovine mastitis, which include inflammation and reddening of the udder tissues, are a host response to bacterial invasion of the mammary gland. In brief, bacterial invasion of the mammary gland triggers the recruitment of neutrophils and expansion of proinflammatory cytokines to control the spread of infection (Boulanger et al., 2003; Persson Waller et al., 2003). This host immune response leads to an increase in the abundance of somatic cells (*i.e.*, neutrophils) in the udder tissue, which can then shed into produced milk (Zhao & Lacasse, 2008; Kehrli & Shuster, 1994). Infections can result from either direct contact with contaminated milking equipment or through environmental exposure post-milking. Contagious, mastitis-causing bacterial pathogens, including host-adapted strains of *Staphylococcus aureus* and *Streptococcus agalactiae*, are primarily spread from cow to cow during the milking process (Jones & Bailey, 2009; Bogni et al., 2011). Environmental pathogens, in contrast, cause opportunistic infections through contact with soiled bedding, manure, or other contaminated areas. A diverse array of environmental bacteria can induce clinical and subclinical forms of mastitis; however, strains of Enterobacteriaceae, Staphylococcaceae, and Streptococcaceae are among the most prevalent causative agents of intramammary infections and are of particular concern to the dairy industry (Jones & Bailey, 2009; Bogni et al., 2011). Historically, most research to control mastitis has focused on disease treatment and intervention of pathogens. Indeed, while much is known about the molecular mechanisms underlying the pathogenesis of mastitis in cows, relatively little is known about the epidemiology of mastitis-causing bacteria in dairy barn environments.

Members of the insect family Muscidae (Insecta: Diptera), including flies of the genera *Musca* (house and face flies) and *Stomoxys* (stable flies), have long been implicated as potential vectors of bovine mastitis (Hansens, 1963; Black & Krafsur, 1985; Semakula et al., 1989; Marley et al., 1991; Meyer & Petersen, 1983). Adult muscid flies are covered in hair-like projections, which may lead to external dissemination of environmental microbes collected from manure or other debris (Schmidt & Roberts, 2013). Dissemination of internally associated bacteria can also occur, either through defecation or regurgitation of saliva when feeding (Sasaki et al., 2000; Butler et al., 1977). Muscid flies are most prevalent on dairy farms during the warmest months of the year, and incidences of mastitis in dairy cattle peak in this same timeframe (Madsen et al., 1991; Smith et al., 1985). *Stomoxys* flies are also obligate blood-feeders, and the dense livestock populations present in most barns provide ample access to nutritional blood meals for biting flies (Meyer & Petersen, 1983). Large fly populations are additionally sustained by the constant availability of raw manure and soiled bedding, which serve as their preferred breeding sites and are also likely to harbor environmental pathogens. In a typical commercial dairy farm setting, adult female flies deposit eggs on both managed manure piles and fresh pats or slurries of manure, where larvae develop through a series of three progressively larger immature stages known as instars before undergoing metamorphosis to the adult stage (Denlinger & Zdárek, 1994). In addition to providing a source of undigested carbohydrates, proteins, and other nutrients to developing larvae, both adult male and female flies are strongly attracted to and may opportunistically ingest cattle manure to support their breeding activities. (Tangtrakulwanich et al., 2015; Albuquerque & Zurek, 2014).

Although historical studies have provided circumstantial evidence linking *Stomoxys* flies to the carriage of mastitis pathogens, very little is known about the native microbiota of biting muscid flies. To date, only a handful of studies have examined the *Stomoxys* microbiota through culture dependent methodology and report the presence of clinically relevant bacterial taxa including, but not limited to, strains of Staphylococcaceae, Enterococcaceae, and Enterobacterales bacteria (Castro et al., 2007; Castro et al., 2010; Schwarz et al., 2020). However, the overall composition and diversity of bacterial communities associated with *Stomoxys* flies, especially in relation to manure habitats, remains to be investigated. In this study, we provide the first culture independent examination of the *Stomoxys* microbiota through longitudinal sampling of adult *Stomoxys* flies and environmental manure collected across two connected dairy facilities. High throughput 16S rRNA gene amplicon sequencing was used to characterize bacterial communities in fly and manure samples collected on a weekly basis during peak fly season in both facilities. Culture dependent methods were then employed to verify the viability of highly prevalent taxa, including mastitis-associated *E. coli*, *Staphylococcus*, and *Klebsiella* spp., in fly homogenates plated on selective enrichment media.

## RESULTS

### *Stomoxys* flies harbor a distinct, low complexity microbiota

*Stomoxys* flies and manure samples were collected on a weekly basis across two dairy facilities in South Central Wisconsin: the UW-Madison Dairy Cattle Center (DCC) and the Arlington Agricultural Research Station Blaine Dairy Cattle Center (Arlington) (Fig. 1). Flies recovered from the same trap were pooled into groups of up to 25 individuals to collect externally associated bacteria before homogenizing subgroups of 3-5 individuals to collect internal bacteria. A total of 183 *Stomoxys* pools (697 flies total) and 106 manure samples were processed for high throughput 16S rRNA gene amplicon sequencing, of which 126 internal fly samples, 32 external fly samples, and 106 manure samples, with a respective median read count of 7108, 5807, and 26050, respectively, were selected for further analysis (Table S1). Across both facilities, bacterial richness was significantly greater in manure samples compared to both internal and external fly samples (Fig. 2), irrespective of sampling date (Fig. S1). Bacterial richness in Arlington-derived external fly samples was also higher than internal fly samples (Fig. 2); however, no significant differences were observed between external and internal fly samples derived from the DCC (Fig. 2; Fig. S1). No significant differences in bacterial richness were observed between manure and fly samples collected from different sampling locations within either facility (Fig. S2), although richness estimates for DCC-derived manure samples were consistently higher than those for Arlington-derived manure samples (Fig. 2; Fig. S1; Fig. S2).

**Fig 1.**
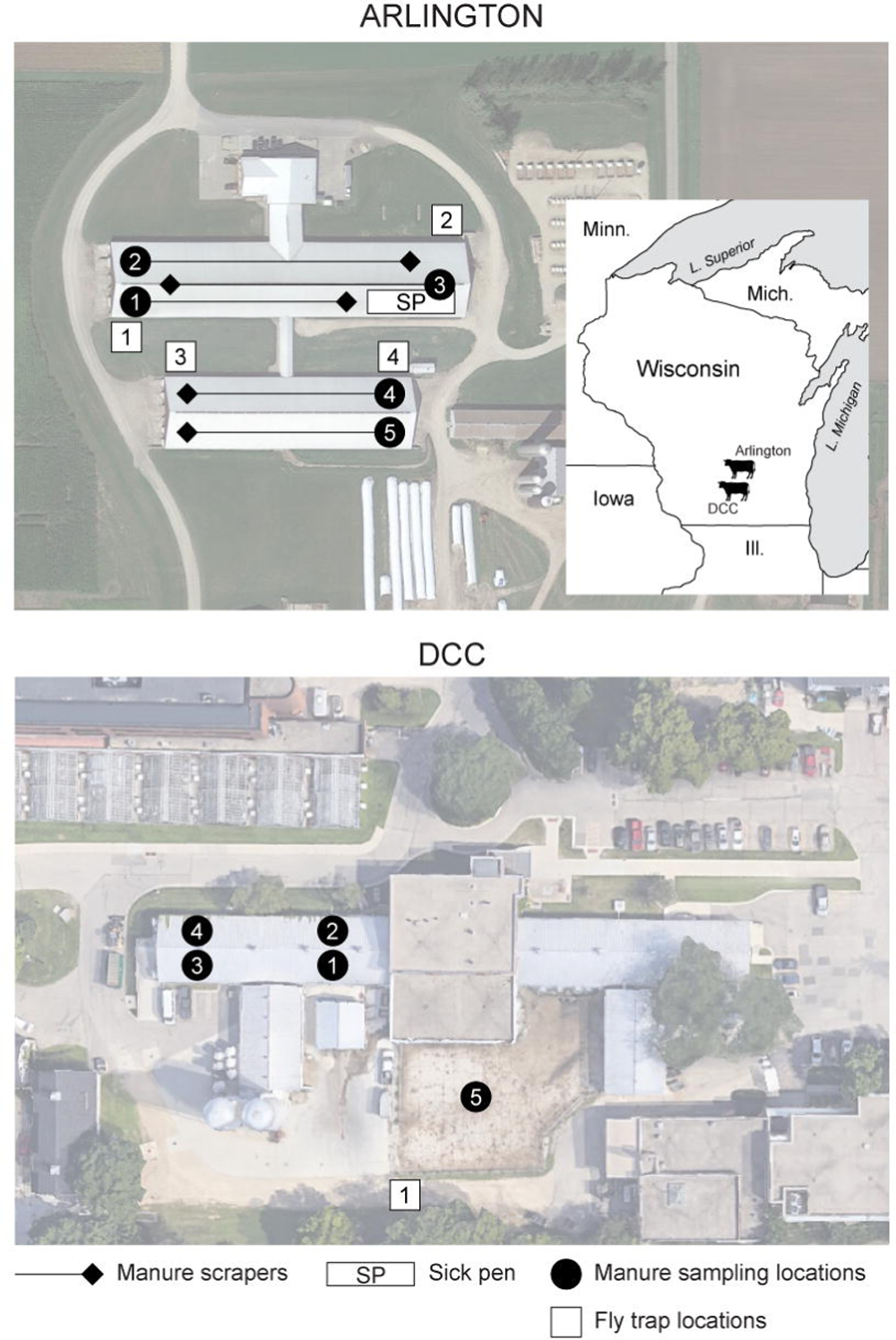
Collection sites for the 16S rRNA libraries and glycerol stocks prepared from manure and fly samples. The upper portion of the figure shows the locations of the collection sites at the Arlington Agricultural Research Station Blaine Dairy Cattle Center (Arlington), while the lower portion of the figure shows locations of the collection sites at the UW-Madison Dairy Cattle Center (DCC). White squares and black circles delineate fly trap and manure sampling locations in both facilities, respectively. The locations of the manure scraper systems and pen housing sick cattle (“Sick pen”) at Arlington are further identified using black diamonds/lines and a white rectangle, respectively (*top panel only*). Both facilities are located in South Central Wisconsin on or in close proximity to the UW-Madison campus.

**Fig 2.**
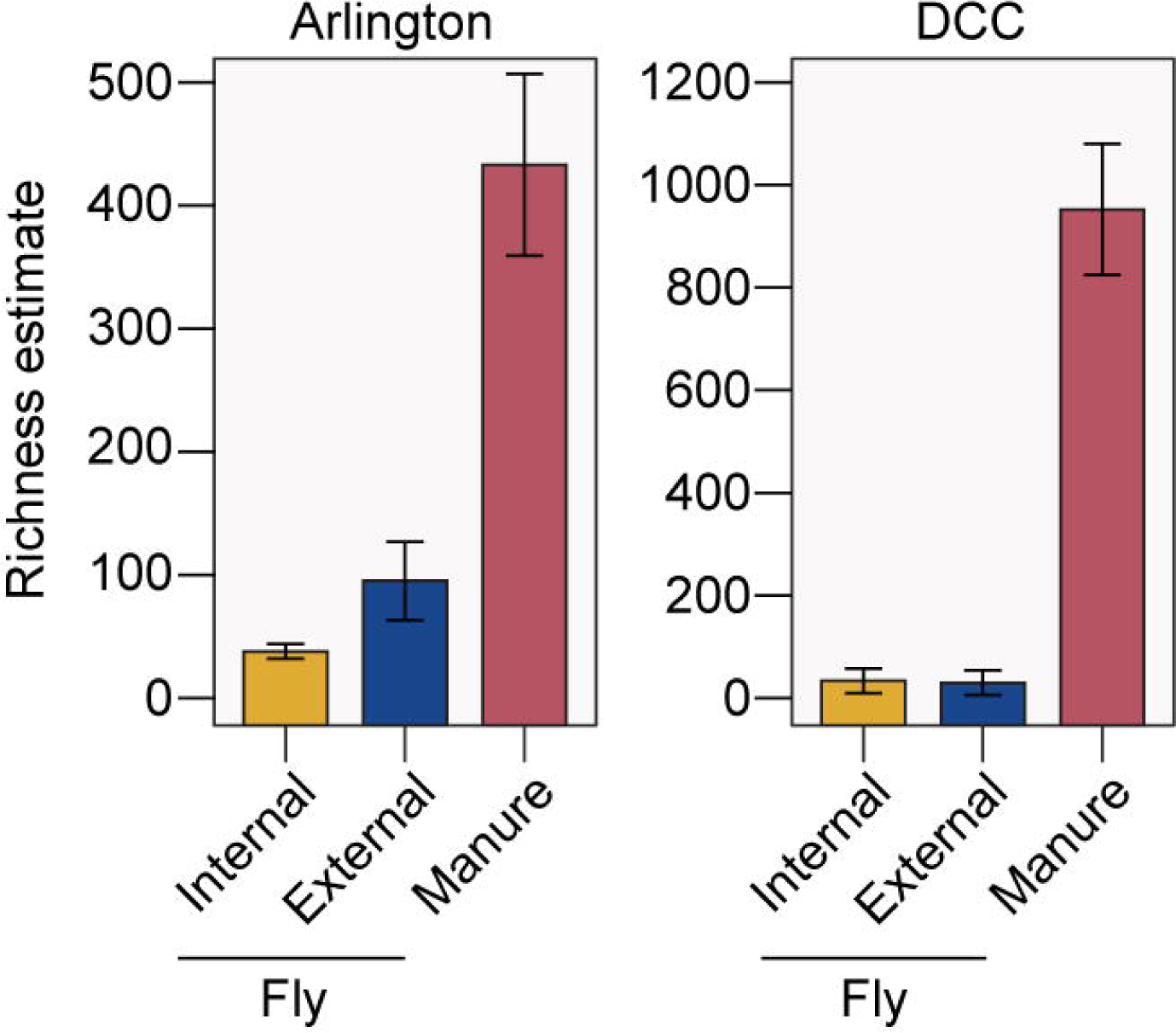
Community-level ASV richness in Arlington- (*left*) and DCC-derived samples (*right*), by sample source (*x*-axis; yellow bars: internal fly samples, blue bars: external fly samples; red bars: manure samples). Bars represent 95% CI (± 1.96 * SE).

Altogether, we identified a total of 7666 bacterial ASVs in manure samples and 1539 and 1233 ASVs across the internal and external fly samples we sequenced, respectively (Table S2). These ASVs corresponded to a total of 191 bacterial orders being represented across all samples, although the relative abundance of each order varied greatly between fly and manure samples (Fig. 3; Table S2). Internal fly samples from both Arlington and the DCC on average harbored high relative abundances of taxa within the Enterobacterales (29.6% Arlington, 37.1% DCC) and Lactobacillales (18.3% Arlington, 20.3% DCC), while taxa within the Staphylococcales were overall more abundant in Arlington internal fly samples (27.7%) than in DCC internal fly samples (11.2%) (ANCOM-BC, log-fold change > 1.5 and FDR-adjusted *p*-value < 0.05) (Fig. 3). Taxa within the Enterobacterales (20.3% Arlington, 27.1% DCC) and Pseudomonadales (11.4% Arlington, 12.1% DCC) were similarly highly abundant in external fly samples from both facilities (Fig. 3). In contrast, manure samples from both Arlington and the DCC on average contained high relative abundances of taxa within the Bacteroidales (15.3% Arlington, 21.9% DCC) and Oscillospirales (21.0% Arlington, 29.4% DCC), while taxa within the Lactobacillales were highly abundant in Arlington manure samples (16.7%) but not DCC manure samples (2.7%) (ANCOM-BC, log-fold change > 1.5 and FDR-adjusted *p*-value < 0.05) (Fig. 3).

**Fig 3.**
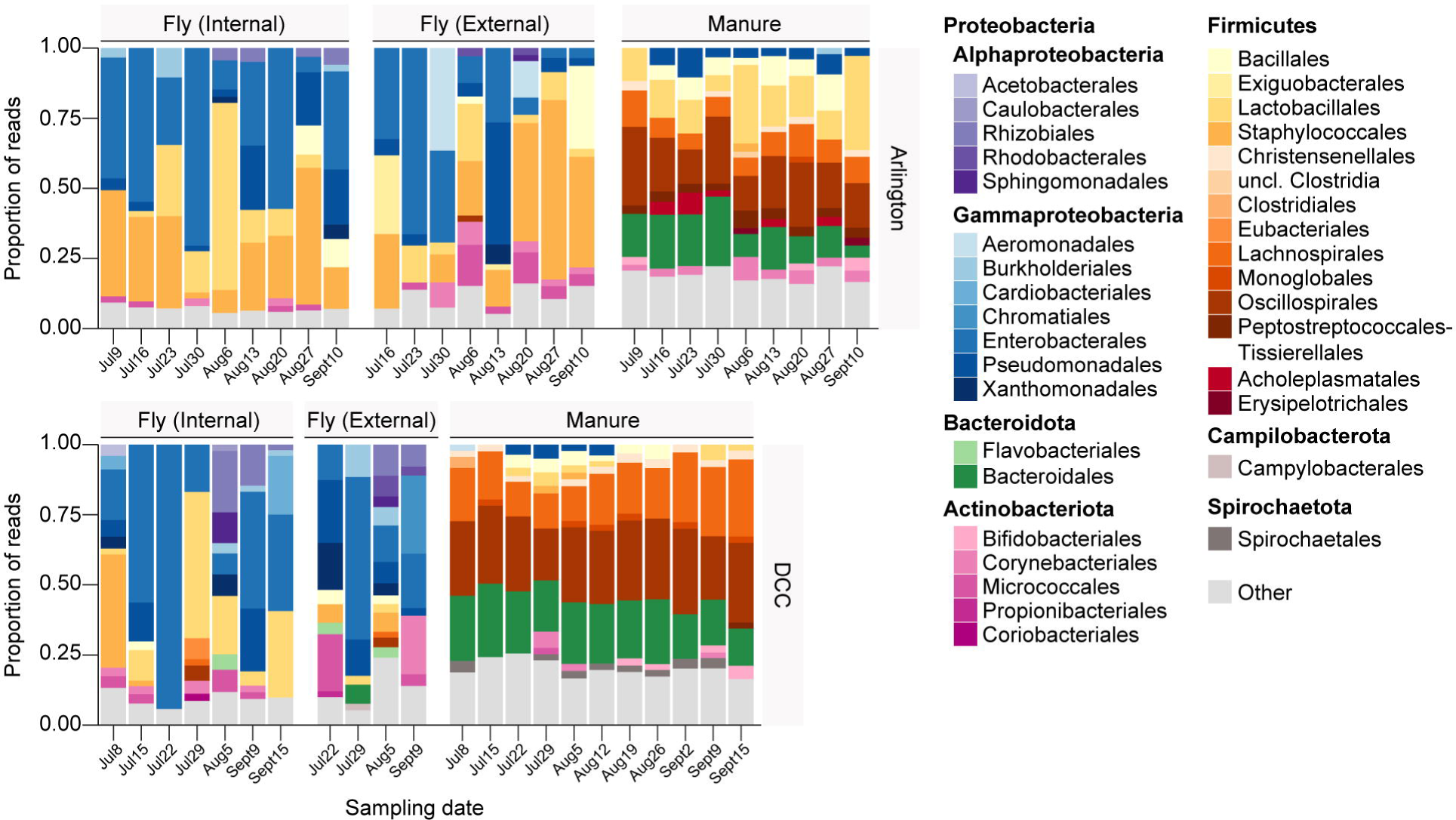
Relative abundance of bacterial orders in Arlington- (*top*) and DCC-derived fly and manure samples (*bottom*), by sampling date (*x*-axis). Libraries derived from the same sample source on a given sampling date were pooled across sampling locations for the bar graphs presented. Colored bars present the proportion of sequencing reads assigned to a given bacterial order. Low abundance orders (<2%) are represented by the “Other” category”.

Principal coordinates analyses using the Bray-Curtis dissimilarity index further revealed significant clustering of bacterial communities by sample source, with both Arlington and DCC manure-associated communities forming distinct clusters separate from bacterial communities in flies collected from the same facilities (Fig. 4), again irrespective of sampling date (Fig. S3; Fig. S4). Arlington-derived internal and external fly samples also formed distinct clusters (Fig. 4), with bacterial communities in Arlington-derived external fly samples generally being more similar to those present in manure collected from the same facility (Fig. 4). In keeping with our bacterial richness results, bacterial community composition was largely consistent across manure and fly samples collected from different sampling locations within a given facility (Fig. S5). The only exceptions to this were the manure samples collected from the sick pen at Arlington, which harbored notably higher relative abundance of bacterial taxa within the order Burkholderiales and lower relative abundances of taxa within the Lactobacillales and Corynebacteriales than manure collected from different sampling locations in the same facility (ANCOM-BC, log-fold change > 1.5 and FDR-adjusted *p*-value < 0.05) (Fig. S5; Fig. S6).

**Fig 4.**
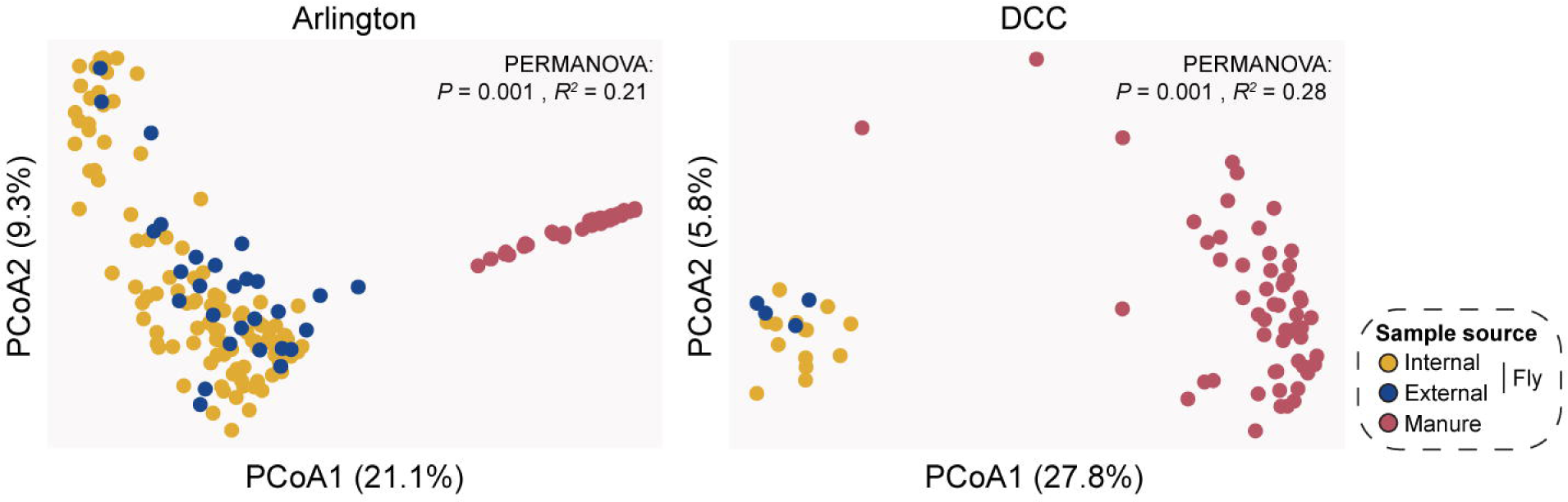
PCoA of Bray-Curtis dissimilarities of community relative abundances, colored by sample source (internal fly samples, yellow; external fly samples, blue; manure samples, red). Each point represents one manure or fly sample community. PERMANOVA identified significant clustering by sample source for both Arlington- (*left*; *P* = 0.001, *R^2^* = 0.21) and DCC-derived samples (*right*; *P* = 0.001, *R^2^* = 0.28). Significant *post hoc* pairwise comparisons between internal/external fly and manure samples were also identified for both facilities (*P* < 0.01). In contrast, a significant pairwise comparison between internal and external fly samples was identified for Arlington (*P* = 0.001) but not the DCC (*P* = 0.2450).

### Bacterial communities in *Stomoxys* flies are highly enriched in mastitis-associated taxa

To develop a more detailed understanding of fly-associated bacterial communities and their relationship to those detected in manure, we next identified bacterial taxa shared between communities in manure and flies collected from the same facility and asked what is the relative abundance of these bacterial taxa in manure and fly samples, respectively. On average, both Arlington and DCC-derived manure and fly samples shared ∼20 bacterial ASVs, which accounted for a large percentage (up to 100%) of the total reads in individual fly samples (Fig. 5), irrespective of sampling date or location (Figs. S7-10). In contrast, these taxa on average represented only ∼39% of the total reads in individual manure samples, although the average relative abundance of shared taxa was significantly lower in manure samples collected from the DCC as compared to those collected from Arlington (Fig. 5; Fig. S11; Fig. S12).

**Fig 5.**
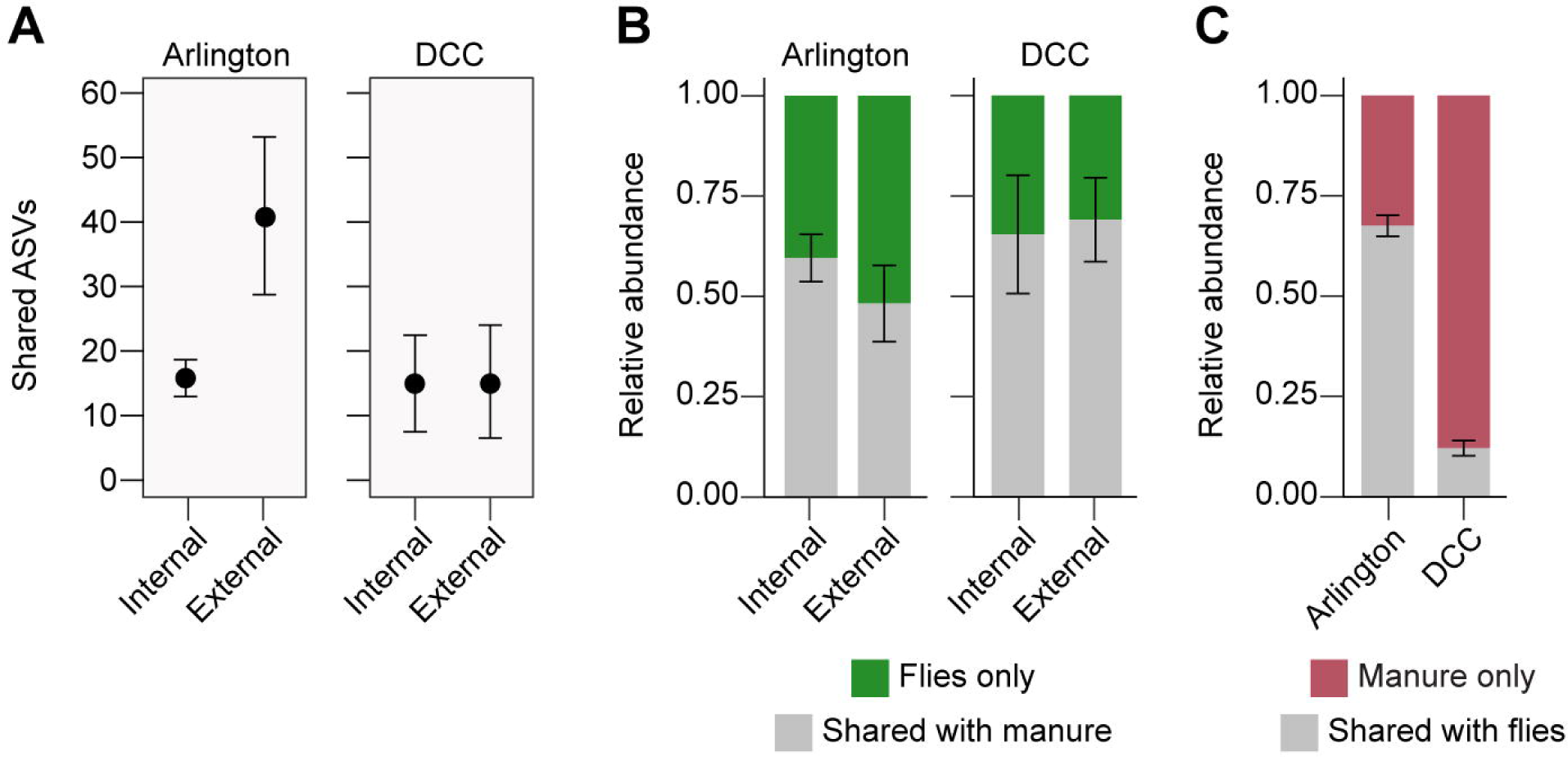
Bacterial ASV transmission analysis. (A) Average number of ASVs shared between Arlington- (*left*) or DCC-derived fly (internal or external) and manure samples (*right*). (B) Average relative abundance of fly associated ASVs (internal or external) shared with manure collected from the same facility. (C) Average relative abundance of manure associated ASVs shared with flies collected from the same facility. Bars in all panels represent 95% CI (± 1.96 * SE).

We also examined whether there were ASVs that were frequently shared by manure and fly samples collected from the same facility. A total of 17 and 12 ASVs were shared in at least 25% of Arlington- and DCC-derived manure-fly sample pairs, respectively (Fig. 6). This included representatives of genera within the bacterial families Enterobacteriaceae (Order Enterobacterales), Enterococcaceae and Streptococcaceae (Order Lactobacillales), Pseudomonadaceae (Order Pseudomonadales), and Staphylococcaceae (Order Bacillales), all of which are commonly associated with environmentally acquired mastitis (Fig. 6) (Klaas & Zadoks, 2018). Perhaps most interesting, however, was the observation that, while the bacterial taxa present in both manure and fly samples represented only a small fraction of manure bacterial communities, they exhibited very high relative abundances in fly bacterial communities (Fig. 7; Figs. S13-16). Highly enriched taxa were also cultured from mastitic cows housed in the same facilities during the sampling period (Table S3).

**Fig 6.**
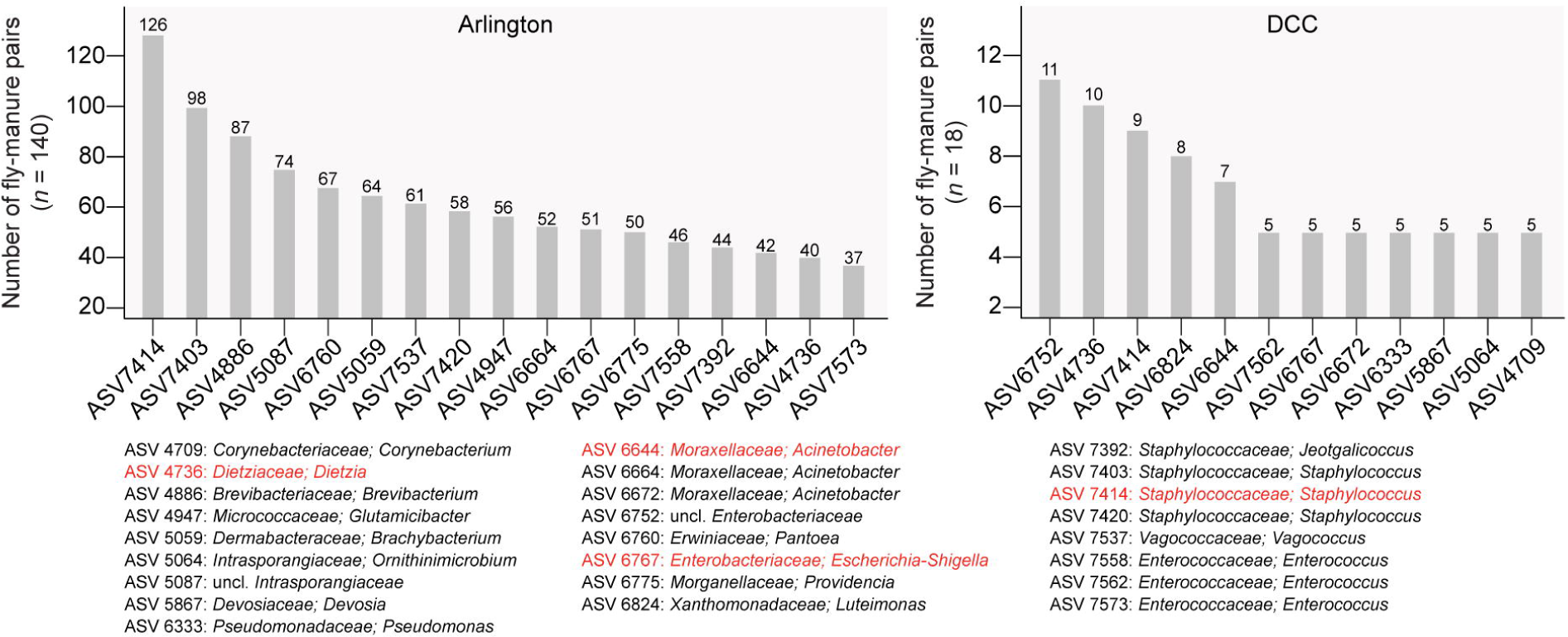
Frequency plot of the 17 and 12 bacterial ASVs shared in at least 25% of Arlington (*left*)- and DCC-derived manure-fly sample pairs (*right*), respectively, along with their taxonomy at the genus level. Taxa highlighted in red were identified as commonly shared ASVs in both facilities.

**Fig 7.**
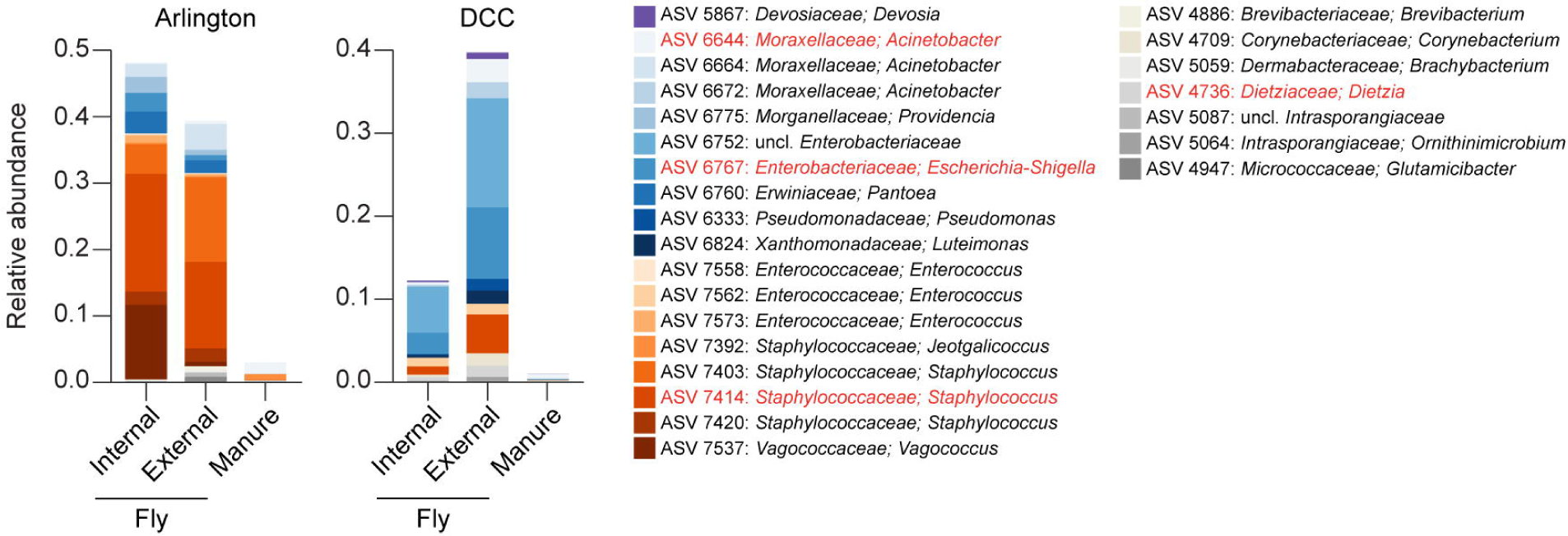
Relative abundance of commonly shared ASVs in Arlington- (*left*) and DCC-derived samples (*right*), by sample source (*x*-axis). Libraries derived from a given sample source were pooled across sampling locations and dates for the bar graphs presented. Colored bars present the proportion of sequencing reads assigned to a given ASV. ASV taxonomy is shown at the genus level. Taxa highlighted in red were identified as commonly shared ASVs in both facilities.

### *Stomoxys* flies serve as *bona fide* reservoirs of viable mastitis-associated bacteria

To confirm that flies carry mastitis-associated bacterial taxa, we screened glycerol stocks generated from homogenates of pooled internal fly samples for the growth of *Staphylococcus, Escherichia*, and *Klebsiella* bacteria. Viable colonies were recovered from 88.5% and 100% of fly homogenates streaked onto MacConkey and Mannitol Salt agar plates, which are generally permissive for the growth of Gram-negative and Gram-positive bacteria, respectively (Table S4). Restreaking of individual colonies on selective agar (followed by morphological or molecular identification) confirmed the presence of *E. coli* in 22.95% (28 of 122) of all plated homogenates, which represents viable growth in 55.81% (24 of 43) of samples with corresponding sequence reads assigned to the genus *Escherichia*-*Shigella* (Table S2; Table S4). The same methods also confirmed the presence of *Klebsiella* spp., which were likely defined as unidentified members of the *Enterobacteriaceae* in our sequencing dataset, in 21.31% (26 of 122) of all plated homogenates, which represents 27.47% (25 of 91) of samples with potential corresponding sequence reads (Table S2; Table S4). Similarly, 75.41% (92 of 122) of all plated homogenates, representing 79.28% (88 of 111) of samples with corresponding sequence reads, contained viable colonies of *Staphylococcus* spp. (Table S2; Table S4).

## DISCUSSION

Muscid flies have long been implicated in the mechanical transmission of environmental bacterial pathogens responsible for mastitis and other bovine diseases (Baldacchino et al., 2013; Mramba et al., 2007). High fly populations are sustained by the large availability of raw manure, which serves as both the preferred oviposition site for adult flies and as a major reservoir of microbial pathogens (Meyer & Petersen, 1983). However, previous experimental studies on the vectorial capacity and microbial communities of muscid flies have focused primarily on the house fly, *Musca domestica*. This work collectively indicates that bacterial isolates can proliferate within the house fly digestive tract leading to the continual excretion of bacteria via regurgitation and defecation (Sasaki et al., 2000; Mramba et al., 2007; Kobayashi et al., 1999; Joyner et al., 2013; Doud & Zurek, 2012). Across sampling locations, *Staphylococcus* and *Weissella* appear to be highly dominant members of the *M. domestica* microbiome; zoonotic pathogens, including *E. coli* O157, *Salmonella enterica*, and *Campylobacter* are also routinely isolated from house fly samples (Bahrndorff et al., 2017; Bahrndorff et al., 2020; Khamesipour et al., 2018; Gupta et al., 2012; Park et al., 2019; Sudagidan et al., 2022; Alam and Zurek, 2004). In contrast, far less is known about the microbiota of biting stable flies. To date, only a handful of studies have used culture dependent methods to survey *Stomoxys* fly-associated microbial communities, identifying strains of *Bacillus*, *Staphylococcus*, and Enterobacterales as among the most common bacterial taxa isolated from adult flies (Castro et al., 2007; Castro et al., 2010; Schwarz et al., 2020). Prior research on *Stomoxys* fly microbiota has also focused exclusively on fly-associated microbes, without also examining potential transmission from the surrounding barn environment or environmental breeding sites.

The first goal of this study was to characterize the *Stomoxys* fly microbiota in relation to environmental manure collected longitudinally over a three-month sampling period across two dairy barns in South Central Wisconsin. Our analyses revealed that adult *Stomoxys* flies harbor a low complexity microbiota dominated by relatively few taxa, which included members of the Enterobacterales, Lactobacillales, and Staphylococcales. In contrast, manure samples harbored much more diverse bacterial communities dominated by taxa within the Bacteroidales, Eubacteriales, and Oscillospirales. Beta diversity analyses likewise revealed that *Stomoxys* bacterial communities clustered separately from manure samples, suggesting some degree of environmental filtering of microorganisms in the fly. Similar patterns of microbial diversity, including both the divergence of host-associated bacterial communities from environmental food sources as well as the relative dominance of specific taxa across fly samples, have also been observed in other terrestrial dipteran species (Bahrndorff et al., 2017; Chandler et al., 2011; Martinson et al., 2017; Martinson et al., 2017b; Jones et al., 2013).

We found that sequenced environmental and *Stomoxys* fly bacterial communities were largely similar across trap locations and collection dates for both facilities; however, there were observable differences in the relative abundances of bacterial taxa across sites. This variability could potentially be explained by either geographical location or differences in sample sizes due to the DCC being a smaller facility in an urbanized area. Cows housed at the DCC are also kept in individual tie-stalls requiring manual manure collection, while Arlington utilizes a free-stall barn system where manure is collected on motorized scraper systems. Cattle and waste management practices impact manure microbiota, which could explain observed differences between bacterial communities in Arlington and DCC manure samples (Tiquia, 2005; Bewley et al., 2017; Hagey et al., 2019; Pandey et al., 2018). Within Arlington manure samples, we also observed a significant decrease in the abundance of Lactobacillales and Corynebacteriales from samples collected within the sick pen. Manure is not collected by a bulk scraper system in the sick pen, and likely represented more recently deposited fecal samples. Differences in diversity could alternatively be attributed to host infection status, which can lead to dysbiosis and increased shedding of microbes in the gastrointestinal tract (Munoz et al., 2006; Ma et al., 2016; Muñoz-Vargas et al., 2018; Pham and Lawley, 2014).

Despite high level taxonomic differences, we still identified specific bacterial ASVs commonly shared between corresponding fly and manure samples. This overlap indicates the potential transmission of a bacterial strain between a manure reservoir and flies. Future studies utilizing metagenomics or whole genome sequencing could both validate the potential of bacterial dissemination in dairy barn environments and help elucidate the genetic diversity of fly-associated bacterial strains. Notably, shared ASVs contributed to a significant portion of reads in the fly microbiome but were also found in relatively low abundance across manure samples, suggesting a possible enrichment of specific manure-associated bacterial strains in the muscid fly gut or a dissemination of fly-associated microbes to the surrounding environment. Our current study did not address whether these taxa were acquired transstadially or via adult feeding activities. In addition to manure, *Stomoxys* flies could alternatively acquire microbes through opportunistic feeding on plant material (Taylor and Berkebile, 2014) or from contact with cattle skin during blood feeding. Laboratory studies *of M. domestica* also suggest that a subset of bacteria is transmitted transstadially from larval to adult flies (Zurek and Nayduch, 2016), although the contribution of transstadial transmission versus ingestion of microbes in shaping the adult fly gut microbiota remains to be elucidated across the Muscidae.

Shared ASVs enriched in fly microbial communities included multiple taxa (*Staphylococcus, Pseudomonas, Enterococcus,* Enterobacterales) associated with environmental bovine mastitis. The high abundance of mastitis associated lineages could be driven by physiological conditions within the dipteran digestive tract (pH, nutrient availability, etc.) which act as a strong selective pressure on microbial community assembly (Engel and Moran, 2013, Lemaitre and Miguel-Aliaga, 2013). Spatial organization of the digestive tract, including colonization of the crop organ, may also allow for the persistence and proliferation of ingested microbes, as has been previously reported in *M. domestica* and *Drosophila* (Sasaki et al., 2000; Pais et al., 2018). In mosquitoes (Diptera: Culicidae), hematophagous activities result in both a decrease in microbial diversity and the selective enrichment of Enterobacteriaceae (Wang et al., 2011; Muturi et al., 2019). Similar selective processes likely impact the *Stomoxys* fly microbiota and could explain the high dominance of Enterobacteriaceae in collected fly samples. In the present study, concurrent farm-specific mastitis incidence records showed that *Staphylococcus* and Enterobacteriaceae are routinely isolated from mastitic cows. Although our study could not address whether fly associated strains represent isolates from clinical cases, the enrichment of bacterial lineages associated with bovine mastitis in the *Stomoxys* microbiota strongly suggests that flies may act as bonafide reservoirs of opportunistic pathogens. Interestingly, we found *Streptococcus* spp., which constitute important contagious and environmental mastitis pathogens, to be rare in *Stomoxys* flies sequenced from both Arlington and the DCC. These results could suggest the possibility that *Streptococcus* species do not efficiently colonize the fly gut. Our results are the first high throughput sequencing of the *Stomoxys* microbiota; further sequencing of *Stomoxys* communities could reveal shifts in taxonomic abundance of mastitis pathogens across geographic regions or the potential of enrichment of specific pathogens during mastitis outbreaks.

The second goal of this study was to verify that fly-associated taxa represent viable and culturable bacterial isolates. In flies, enzymatic activity and other digestive processes can facilitate the lysis of bacteria within the gut (Ludington and Ja, 2020; Nayduch and Burrus, 2017). The use of 16S rRNA sequencing data has been critical to our understanding of animal microbiomes; however, amplification of DNA can be from either viable or dead cells within samples (Young et al., 2007; Cangelosi and Meschke, 2014). Fly pools were therefore screened for culturable *Klebsiella*, *E. coli*, and *Staphylococcus* spp., which represent key mastitis-associated bacterial pathogens from highly dominant lineages across fly samples. Our results show that taxa associated with mastitis were not only observed in the sequence data but could be readily cultured from a subset of the corresponding fly homogenates. Together, these results confirm that *Staphylococcus* and Enterobacteriaceae are prominent members of the *Stomoxys* microbiota. The culture-dependent methodology is likely an underrepresentation of the true incidence rates as culturing was performed on an enriched aliquot of each fly homogenate pool. We also observed culturable isolates in a subsection of samples without corresponding sequence reads, which could indicate those taxa were rare in the unenriched homogenates used for DNA extraction and amplicon sequencing.

Collectively, our results provide evidence for a potential role for *Stomoxys* flies in the carriage of opportunistic pathogens in dairy barns. Our approach, which included parallel collections of flies and associated environmental manure samples, represents the first high throughput 16S rRNA amplicon sequencing of the *Stomoxys* fly microbiota. We found that mastitis-associated bacterial taxa were highly dominant across fly samples, but relatively rare across corresponding manure samples. Viable colonies of mastitis associated taxa were also readily isolated from fly samples, indicating that the *Stomoxys* fly may represent an important, yet understudied, element of disease transmission in agricultural settings. Future work will elucidate the role of biting flies in the transmission of not only bovine mastitis pathogens, but other farm-associated zoonotic pathogens including *E. coli* O157, *Brucella*, and *Salmonella* (Ruegg et al., 2003; Seleem et al., 2010). Future work will also leverage these results to provide novel insights into the ecology and evolution of these and other dipteran insects more broadly, including species that serve important roles as bioindicators, biocontrol agents, sources of nutrition for other organisms, and/or nuisance pests, as well as other vectors of disease (Arellano et al., 2023).

## MATERIALS AND METHODS

### Field sampling

Samples of *Stomoxys* flies and manure were collected on a weekly basis from July through September 2021 across two focal dairy farms in South Central Wisconsin: the UW-Madison Dairy Cattle Center (DCC) and the Arlington Agricultural Research Station Blaine Dairy Cattle Center (Arlington) (Fig. 1). *Stomoxys* flies were caught on adhesive alsynite fiberglass traps (Olson Products, Medina, OH, USA), which selectively attract *Stomoxys* flies through reflection of ultraviolet light (Agee & Patterson, 1983; Hogsette & Ruff, 1990). A single trap was set adjacent to an outdoor cow pen at the DCC, while four fly traps were set at Arlington outside the free-stall barn structures (Fig. 1). Traps at both farms were retrieved and replaced weekly. For manure samples, at least 25 ml of material was collected in a sterile conical tube for downstream processing (Thermo Fisher Scientific, Waltham, MA, USA). A total of five manure samples were collected each week from the DCC–one pooled from material collected along each quadrant of the associated tie-stall barn and one containing material collected from an outdoor pen (Fig. 1). Six manure samples were collected each week from Arlington–one pooled from material collected from each of five manure scraper systems and one containing material from an isolated pen housing sick cattle in the same facility (Fig. 1). Upon retrieval, all samples were transported directly to the laboratory on ice. Fly samples were immediately stored at −20 °C until processing; manure samples were diluted 1:1 with sterile 1X phosphate-buffered saline (PBS) and vortexed to ensure homogenization before storing at −80 °C (Carroll et al., 2012).

Biting flies belonging to the genus *Stomoxys* were identified with the assistance of taxonomic keys available in the *Manual of Nearctic Diptera* (McAlpine et al., 1987). Whole bodies of flies from each trap were carefully retrieved from adhesive linings with surface-sterilized featherweight tweezers and pooled together into groups of up to 25 individuals. Pools were then vortexed gently for 40 sec in 10 ml PBS-T (1X PBS + 0.01% Tween 80), and flies were removed before centrifugation of the PBS-T solutions to isolate the external fly microbiota. To isolate the internal fly microbiota, the same flies were subsequently surface sterilized with successive washes in 70% ethanol, 0.05% bleach, and water, and homogenized in groups of 3-5 flies by bead-beating with 3 x 5 mm stainless steel beads (Qiagen, Hilden, Germany) in 1 ml 1X PBS. For each homogenate, a 40 μl aliquot was plated onto both Tryptic Soy agar (TSA) and Brain Heart Infusion (BHI) agar plates before incubating at 30 °C for 3 days. The resulting microbial growth was collected in 1 ml sterile 1X PBS and stored as a 40% glycerol stock at −80 °C for use in bacterial isolations. The remainder of each homogenate was then centrifuged at 1,000 rcf (x *g*) to concentrate any remaining insect integument, which could interfere with DNA extractions and PCR amplification (Rubin et al., 2014; Beckmann & Fallon, 2014), and supernatants were transferred to new tubes prior to centrifugation at 21,300 rcf (x *g*) for 20 min to pellet any cells. All cell pellets were stored at −20 °C prior to DNA isolation and sequencing.

### Bacterial 16S rRNA library construction and sequencing

Total genomic DNA was isolated from fly-derived cell pellets, 200 μl of homogenized manure samples, and associated extraction controls using a Qiagen DNeasy Blood & Tissue kit prior to one-step PCR amplification of the V4 region of the 16S rRNA gene using barcoded primers as described previously (Kozich et al., 2013). No-template reactions as well as reactions using templates from blank DNA extractions served as negative controls. With the exception of 16 external and 6 internal fly samples amplified at 35 cycles, PCR amplification was performed in 25 μl reactions under the following conditions: initial denaturation cycle of 95 °C for 3 min, followed by 30 cycles at 95 °C for 30 sec, 55 °C for 30 sec and 72 °C for 30 sec, and a final extension step at 72 °C for 5 min. PCR products were visualized on 1% agarose gels, purified using a MagJET NGS Cleanup and Size Selection Kit (Thermo Fisher Scientific, Waltham, MA, USA), and quantified with a Quantus fluorometer (Promega, Madison, WI, USA). Purified libraries were then combined in equimolar amounts prior to paired-end sequencing (2L×L250 bp) on an Illumina MiSeq at the University of Wisconsin-Madison.

### Sequence processing and data analysis

Paired end demultiplexed sequences were imported into QIIME2 2022.2.0 for processing (Bolyen et al., 2019). Sequence quality scores were assessed and denoising was performed via DADA2 (Callahan et al., 2016), followed by multiple sequence alignment and phylogenetic tree construction using Mafft and FastTree2, respectively (Katoh & Standley, 2013; Price et al., 2010). Taxonomy was assigned using a Naive-Bayes classifier natively implemented in Qiime and pre-trained against the SILVA reference database (Quast et al., 2013). The resulting Qiime2-generated taxonomy table, ASV table, and phylogenetic tree were then imported into R (version 4.1.3) and merged with sample metadata as phyloseq objects (McMurdie & Holmes, 2013). Suspected contaminants were identified and removed from the ASV table using the ‘decontam’ R package, which compares both the frequency and prevalence of ASVs between samples, extraction controls, and PCR controls (Davis et al., 2018). ASV reads classified as “Archaea”, “Chloroplast”, or “Mitochondria”, as well as samples containing fewer than 100 reads, were also removed prior to further analysis.

Data analysis was performed in R version 4.3.1 (http://www.r-project.org/), using “ggplot2” (Wickham, 2016) for data visualization. Bacterial community richness was estimated using the weighted linear regression model of ASV richness estimates to calculate 95% confidence intervals (CIs) for group means using the *betta()* function in “breakaway” (Willis & Bunge, 2022), interpreting only groups with non-overlapping 95% CIs. Beta diversity was visualized for Bray-Curtis dissimilarities (Bray & Curtis, 1957) of relative abundance data using principal coordinates analysis (PCoA) via the *ordinate()* function in “phyloseq” (McMurdie & Holmes, 2013). To test for a significant effect of sampling source or sampling date on community composition, we used permutational multivariate analysis of variance (PERMANOVA) to partition Bray-Curtis dissimilarity matrices among sources of variation using the *adonis()* function in “vegan” (Anderson, 2001) and permutational multivariate analysis of dispersion (PERMDISP) to test for homogeneity of dispersion using the *betadisper()* function (Anderson, 2006). Significant PERMANOVA and PERMDISP results were then subjected to *post hoc* pairwise comparisons, adjusting *p*-values using the Benjamini-Hochberg False Discovery Rate (FDR) method (Benjamini & Hochberg, 1995) to identify significant differences between groups. Finally, the package “ANCOM-BC” (Lin & Peddada, 2020) was used to identify significant changes in taxa abundance between sample groups, interpreting only taxa with an effect size (log-fold change) > 1.5 and FDR-adjusted *p*-value < 0.05 for a given group comparison.

### Isolation of mastitis-associated bacterial taxa

Selective microbial growth media was used to isolate select clinically relevant bacterial taxa from fly-derived glycerol stocks. In brief, stocks were streaked onto MacConkey agar (lactose base) plates to enrich the growth of gram-negative enteric bacteria. Presumptive *Escherichia coli* colonies (*i.e.*, showing both lactose fermentation and bile salt precipitation) were re-streaked onto Eosin Methylene Blue (EMB) agar plates where they were verified as *E. coli* if a distinctive green, metallic sheen was observed (Leininger et al., 2001). *Klebsiella* strains were isolated on MacConkey-inositol-carbenicillin agar, which allows for the enrichment and subsequent differentiation of *Klebsiella* spp. from other *Enterobacteriaceae* (Bagley & Seidler, 1978). Presumptive *Klebsiella* isolates were confirmed by comparison against the NCBI BLAST database after performing Sanger sequencing of the 16S rRNA gene using the universal 1492R primer (Frank et al., 2008). Mannitol Salt agar was used as a primary selective media to enrich the growth of *Staphylococcus* spp. Presumptive *Staphylococcus* species were verified via colony PCR using the *Staphylococcus*-specific primers TStaG422 and TStag765 (Martineau et al., 2001).

## SUPPLEMENTAL MATERIAL

**Supplemental file 1: Table S1** Sample metadata and sequencing statistics.

**Supplemental file 2: Table S2** Prevalence of each ASV in each sample, along with its taxonomic assignment.

**Supplemental file 3: Table S3** Etiology of infection for symptomatic cows housed at Arlington or the DCC during the study sampling period.

**Supplemental file 4: Table S4** Culture results for glycerol stocks generated from internal fly samples, along with whether a given taxon was detected in each sample via high throughput 16S rRNA gene amplicon sequencing.

**Supplemental file 5:** Figures S1-S16.

## DATA AVAILABILITY

Raw Illumina reads are available in the NCBI Sequence Read Archive (https://www.ncbi.nlm.nih.gov/sra) under BioProject ID PRJNA1032128. Scripts used for analysis and figure generation are available in the Coon laboratory’s GitHub repository (https://github.com/kcoonlab/stable-fly-microbiome). All other data generated by this study are available as Supporting Information herein.

## ACKNOWLEDGMENTS

We thank Jessica Cederquist and the UW-Madison Department of Animal and Dairy Science Dairy Herd Operation for access to the dairy cattle facilities included in this study (Arlington and the DCC), along with their associated records. We also thank Garret Suen and Johanna Elfenbein (UW-Madison) for advice on study design and members of the Coon laboratory (Aldo Arellano, Miguel Medina Muñoz, Holly Nichols, and Serena Zhao) for helpful feedback and assistance with field collections. This work was supported by awards from the UW Dairy Innovation Hub (to KLC). AJS was additionally supported by a USDA AFRI EWD Predoctoral Fellowship (2023-67011-40337) and mini grant from the UW Center for Integrated Agricultural Systems. AJS and KLC conceived of and designed the experiments. AJS, JEK, and KLC performed the experiments. AJS and KLC carried out the data analysis. AJS wrote the initial manuscript, and JEK and KLC contributed to revisions.

