## Supplemental Figures for "Stable flies are bonafide reservoirs of mastitis-associated bacteria"

**Supplementary information for:** Stable flies are bonafide reservoirs of mastitis-associated bacteria

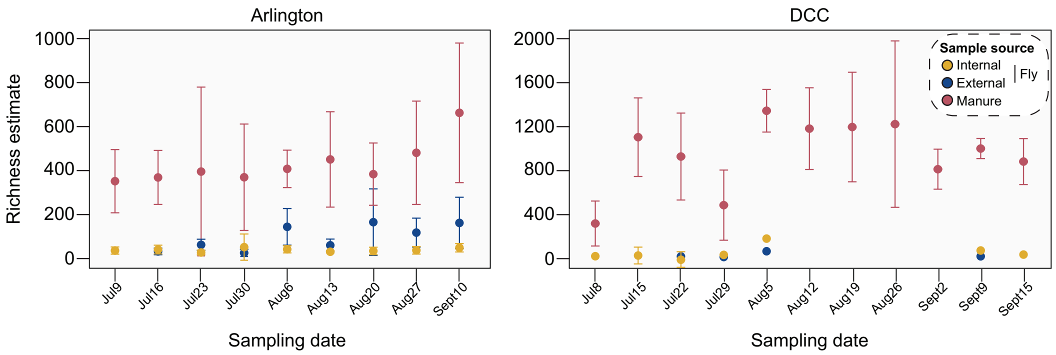


**Fig S1** Community-level ASV richness in Arlington- (*left*) and DCC-derived fly and manure samples (*right*), by sampling date (*x*-axis). Points are colored by sample source (internal fly samples, yellow; external fly samples, blue; manure samples, red). Bars represent 95% CI (± 1.96 * SE).

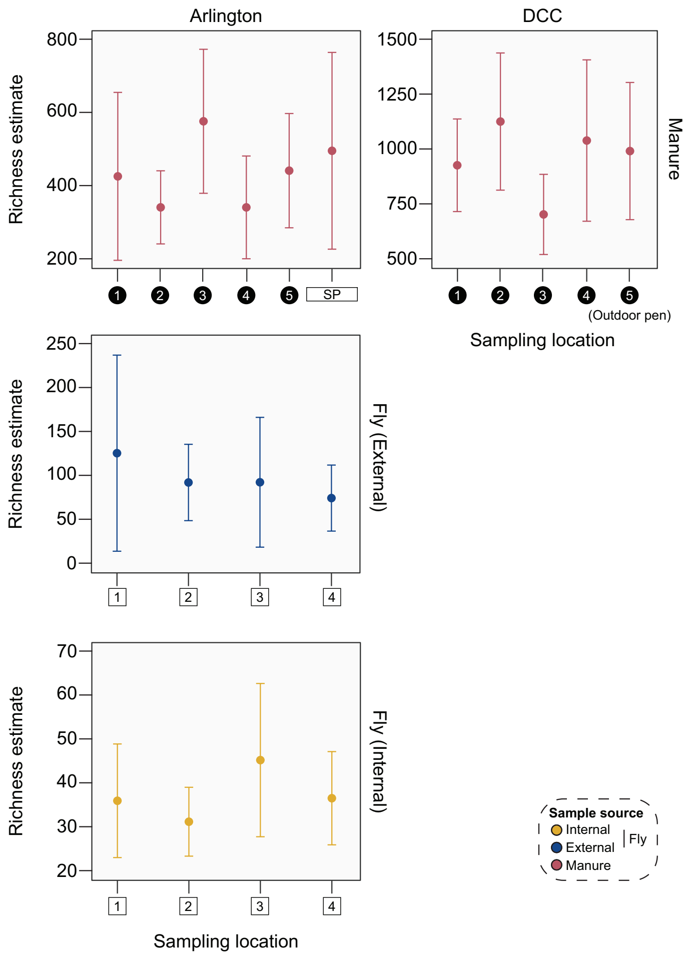


**Fig S2** Community-level ASV richness in Arlington- (*left*) and DCC-derived fly and manure samples (*right*), by sampling location (*x*-axis; see Figure 1 for more information). Points are colored by sample source (top panel: manure samples, red; middle panel: external fly samples, blue; bottom panel: internal fly samples, yellow). Bars represent 95% CI (± 1.96 * SE).

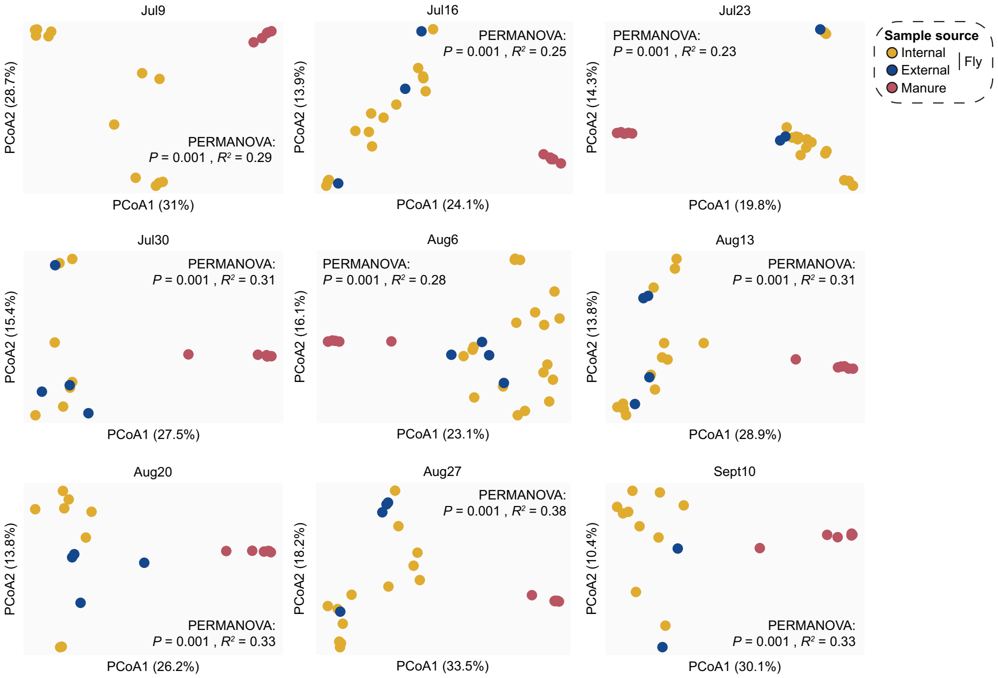


**Fig S3** PCoA of Bray-Curtis dissimilarities of community relative abundances, by sampling date. Points are colored by sample source (internal fly samples, yellow; external fly samples, blue; manure samples, red), with each point representing one Arlington-derived manure or fly sample community. PERMANOVA identified significant clustering by sample source for all sampling dates ($P$ = 0.001, $R^2$ = 0.23-0.38).

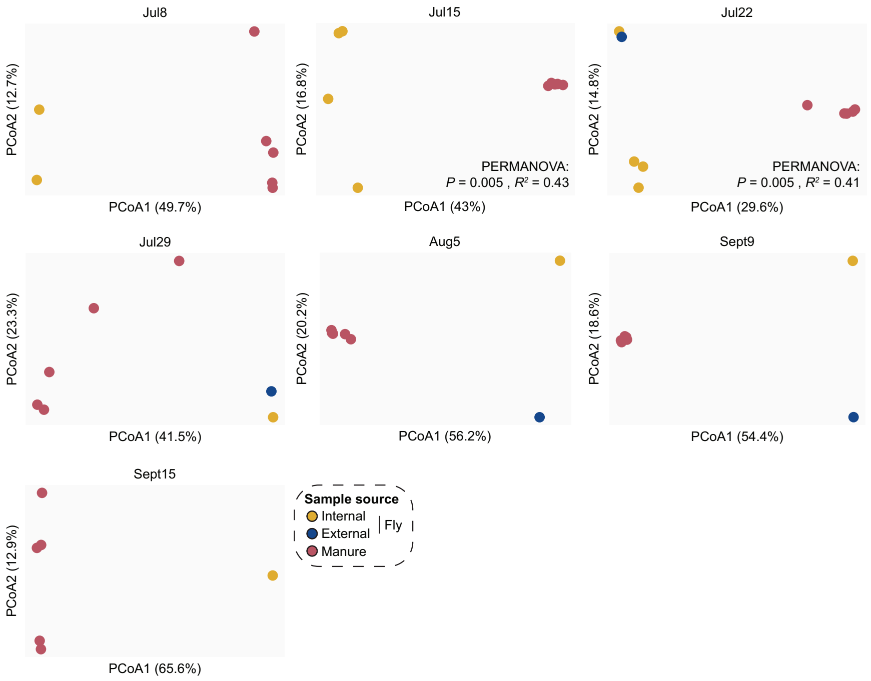


**Fig S4** PCoA of Bray-Curtis dissimilarities of community relative abundances, by sampling date. Points are colored by sample source (internal fly samples, yellow; external fly samples, blue; manure samples, red), with each point representing one DCC-derived manure or fly sample community. PERMANOVA identified significant clustering by sample source for both sampling dates for which there were enough samples to compare communities (Jul15: $P = 0.008$, $R^2 = 0.43$; Jul22: $P = 0.007$, $R^2 = 0.41$).

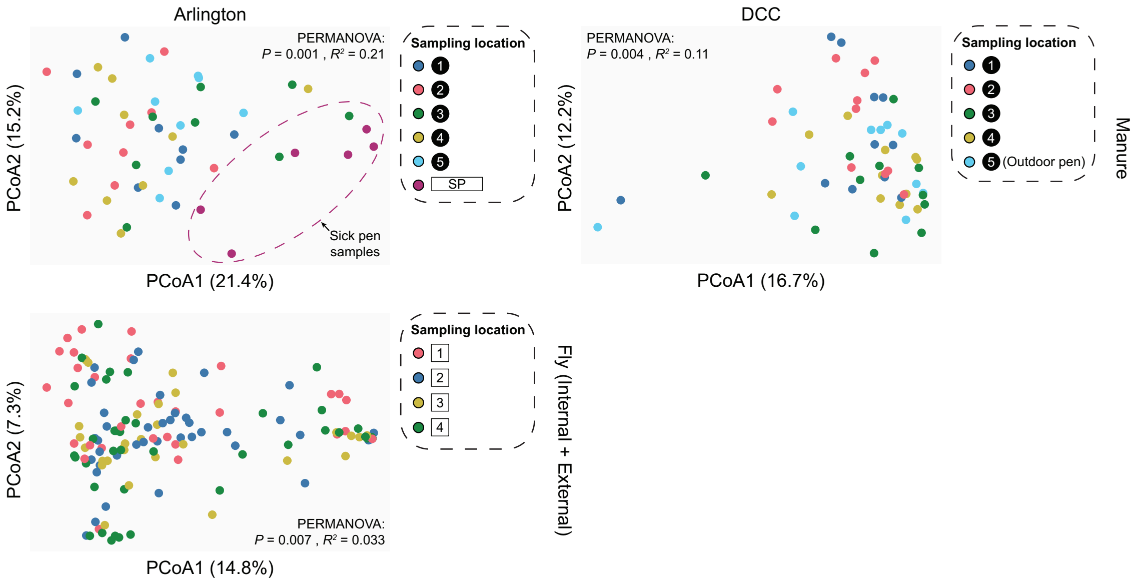


**Fig S5** PCoA of Bray-Curtis dissimilarities of community relative abundances, colored by sampling location (see Figure 1 for more information). Each point represents one Arlington- (*left*) or DCC-derived manure (*top*) or fly *(bottom*) sample community. PERMANOVA identified significant clustering by sampling location for both Arlington- (manure: $P$ = 0.001, $R^2$ = 0.21; fly: $P$ = 0.003, $R^2$ = 0.03) and DCC-derived samples (manure: $P$ = 0.006, $R^2$ = 0.11). However, significant *post hoc* pairwise comparisons were only identified for Arlington- and DCC-derived manure samples, including those between samples collected from the Arlington sick pen and other sampling locations at the same facility ($P$ < 0.05). No significant pairwise comparisons were identified for Arlington-derived fly samples ($P$ > 0.05).

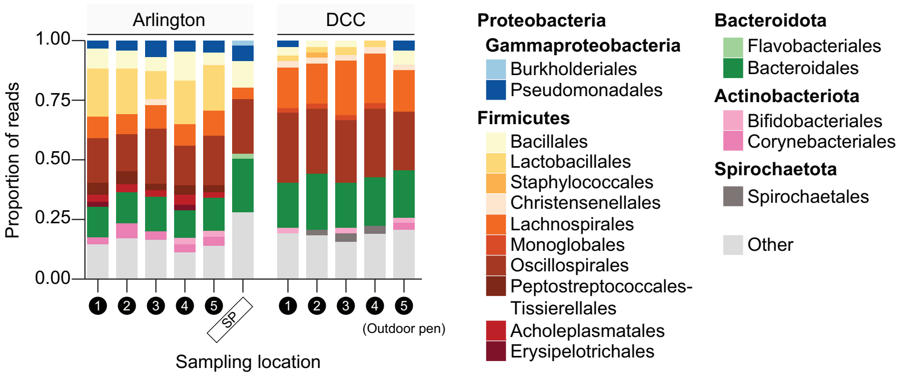


**Fig S6** Relative abundance of bacterial orders in Arlington- (*left*) and DCC-derived manure samples (*right*), by sampling location (*x*-axis; see Figure 1 for more information). Libraries derived from samples collected at a given sampling location were pooled across sampling dates for the bar graphs presented. Colored bars present the proportion of sequencing reads assigned to a given bacterial order. Low abundance orders (<2%) are represented by the "Other" category".

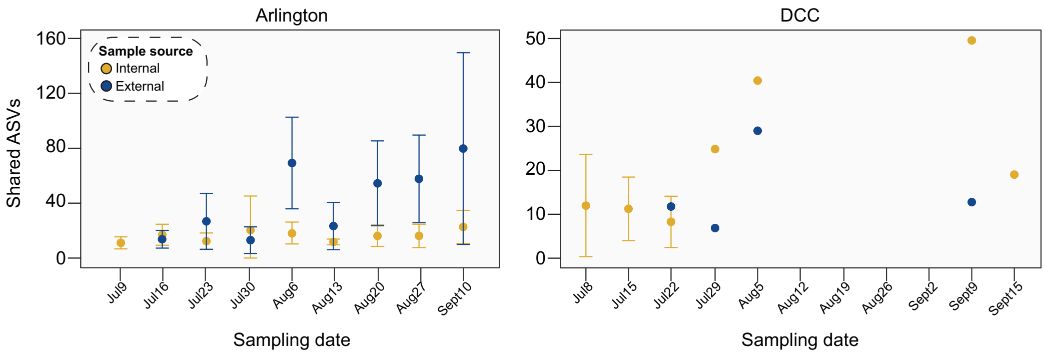


**Fig S7** Average number of ASVs shared between Arlington- (*left*) or DCC-derived fly (internal or external) and manure samples (*right*), by sampling date (*x*-axis). Points are colored by sample source (internal fly samples, yellow; external fly samples, blue). Bars represent 95% CI (± 1.96 * SE).

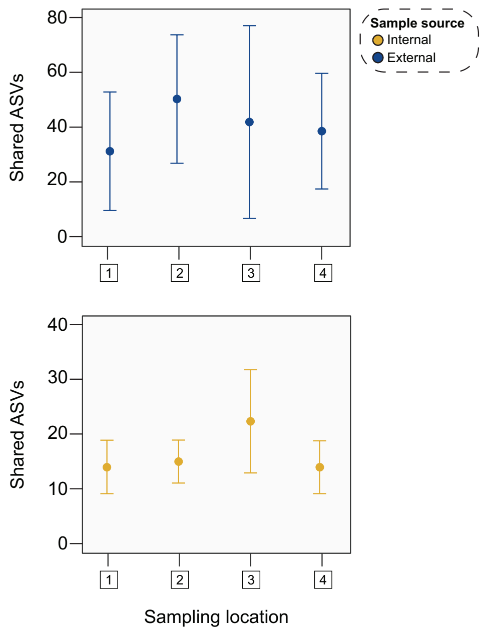


**Fig S8** Average number of ASVs shared between Arlington-derived fly (internal or external) and manure samples, by sampling location (*x*-axis; see Figure 1 for more information). Points are colored by sample source (top panel: external fly samples, blue; bottom panel: internal fly samples, yellow). Bars represent 95% CI (± 1.96 * SE).

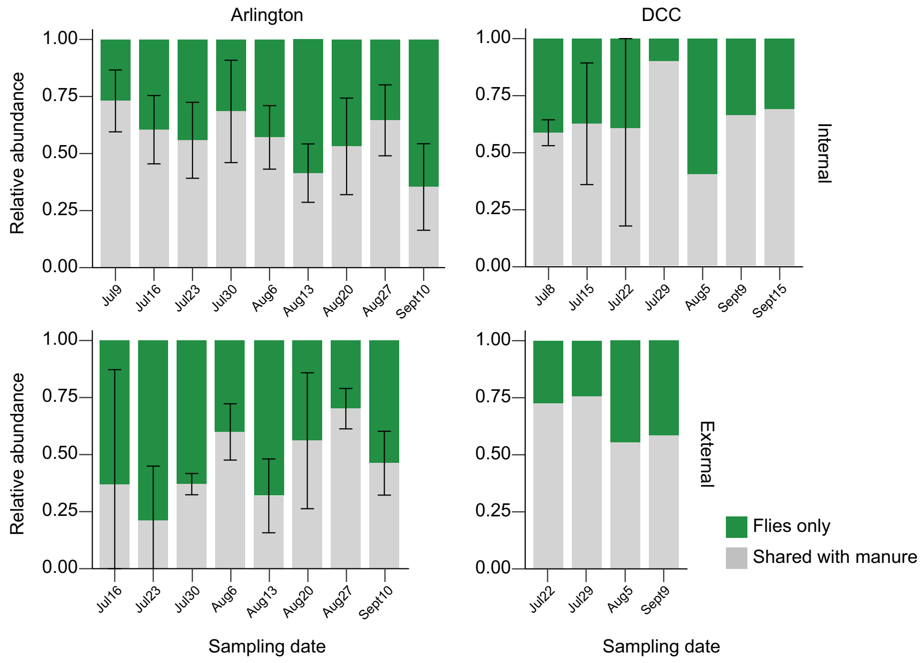


**Fig S9** Average relative abundance of fly-associated ASVs (internal or external) shared with manure collected from Arlington (*left*) or the DCC (*right*), by sampling date (*x*-axis). Bars represent 95% CI (± 1.96 * SE).

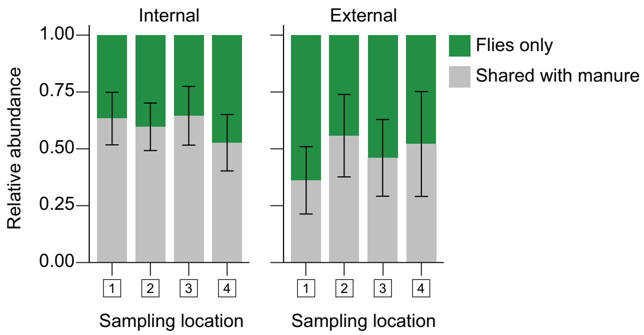


**Fig S10** Average relative abundance of fly-associated ASVs (internal or external) shared with manure collected from Arlington, by sampling location (*x*-axis; see Figure 1 for more information). Bars represent 95% CI (± 1.96 * SE).

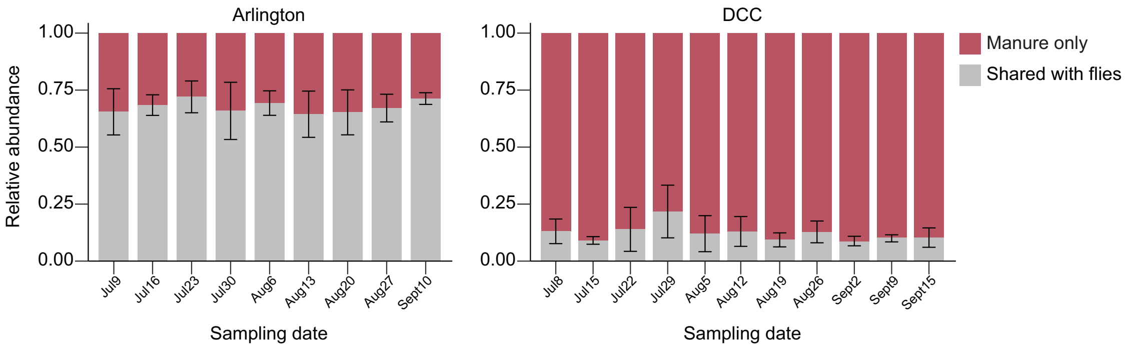


**Fig S11** Average relative abundance of manure-associated ASVs shared with flies collected from Arlington (*left*) or the DCC (*right*), by sampling date (*x*-axis). Bars represent 95% CI (± 1.96 * SE).

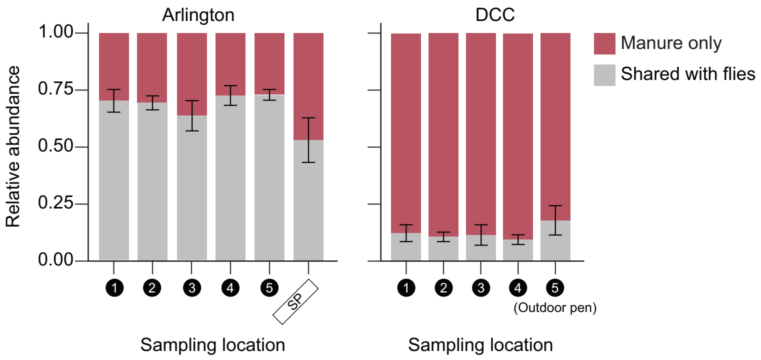


**Fig S12** Average relative abundance of manure-associated ASVs shared with flies collected from Arlington (*left*) or the DCC (*right*), by sampling location (*x*-axis; see Figure 1 for more information). Bars represent 95% CI (± 1.96 * SE).

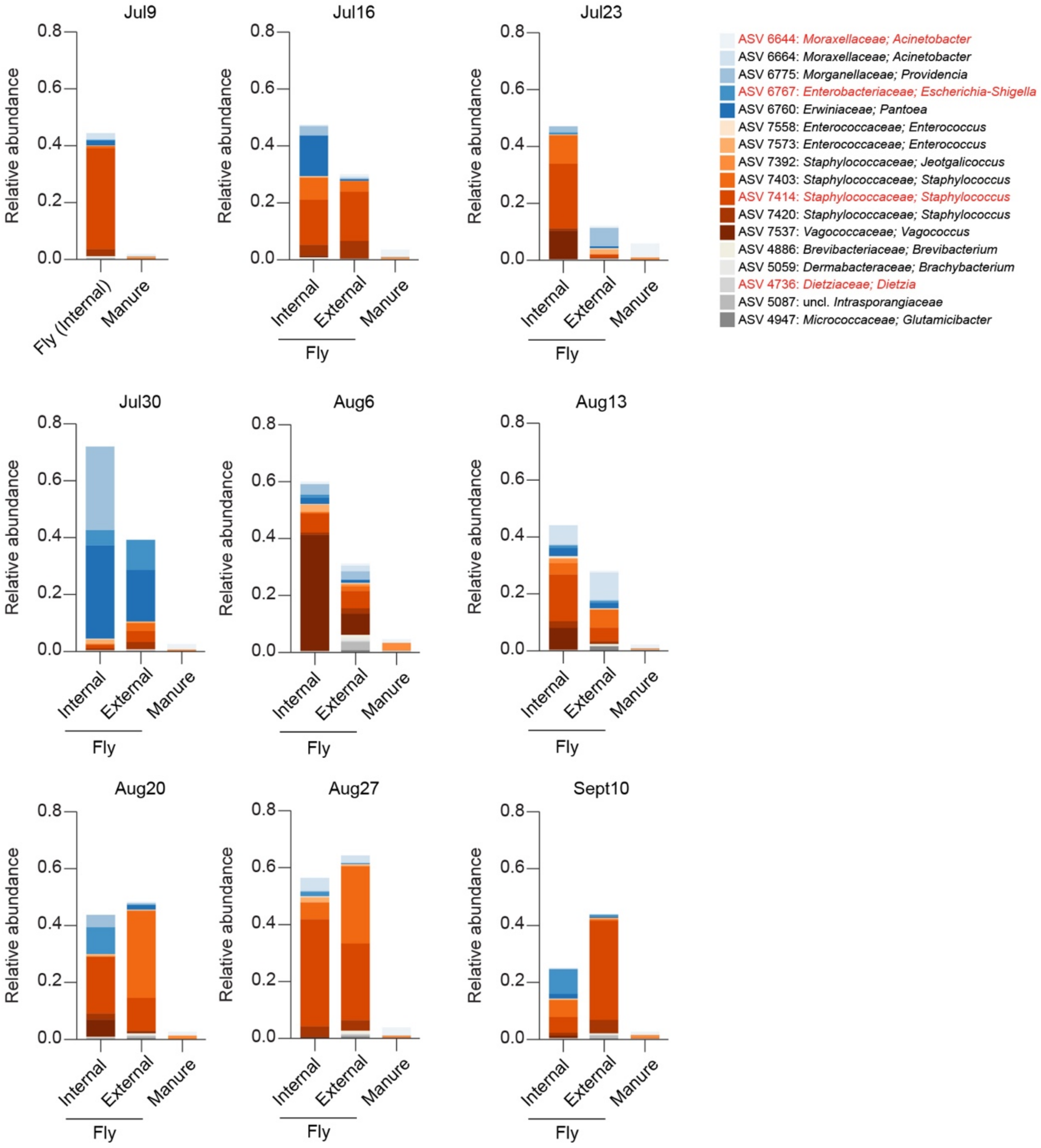


**Fig S13** Relative abundance of commonly shared ASVs in Arlington-derived fly and manure samples, by sampling date. Libraries derived from the same sample source on a given sampling date were pooled across sampling locations for the bar graphs presented. Colored bars present the proportion of sequencing reads assigned to a given ASV. ASV taxonomy is shown at the genus level. Taxa highlighted in red were identified as commonly shared ASVs in both facilities.

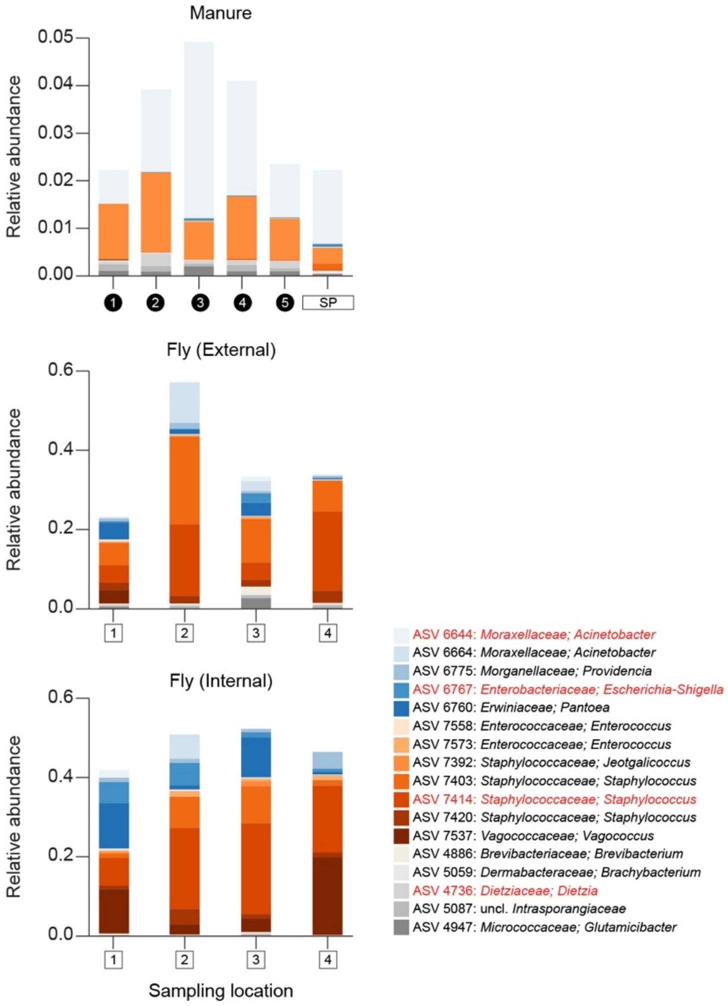


**Fig S14** Relative abundance of commonly shared ASVs in Arlington-derived fly and manure samples, by sampling location (*x*-axis; see Figure 1 for more information). Libraries derived from samples collected from the same sample source at a given sampling location were pooled across sampling dates for the bar graphs presented. Colored bars present the proportion of sequencing reads assigned to a given ASV. ASV taxonomy is shown at the genus level. Taxa highlighted in red were identified as commonly shared ASVs in both facilities.

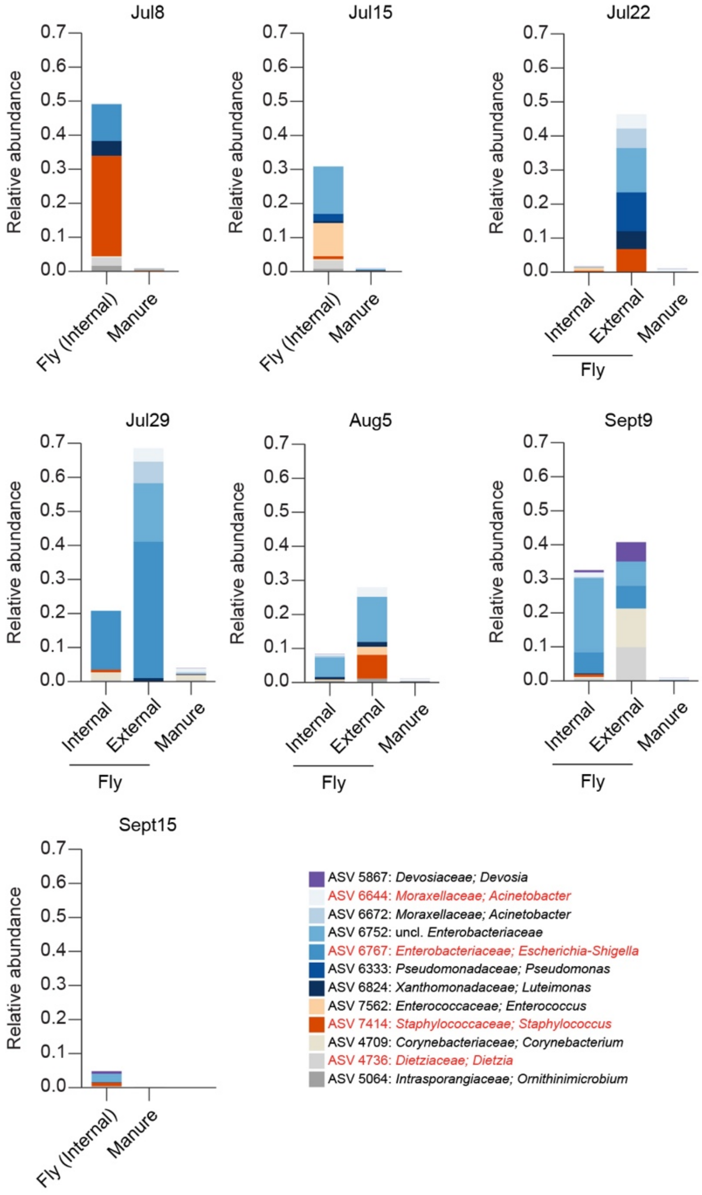


**Fig S15** Relative abundance of commonly shared ASVs in DCC-derived fly and manure samples, by sampling date. Libraries derived from the same sample source on a given sampling date were pooled across sampling locations for the bar graphs presented. Colored bars present the proportion of sequencing reads assigned to a given ASV. ASV taxonomy is shown at the genus level. Taxa highlighted in red were identified as commonly shared ASVs in both facilities.

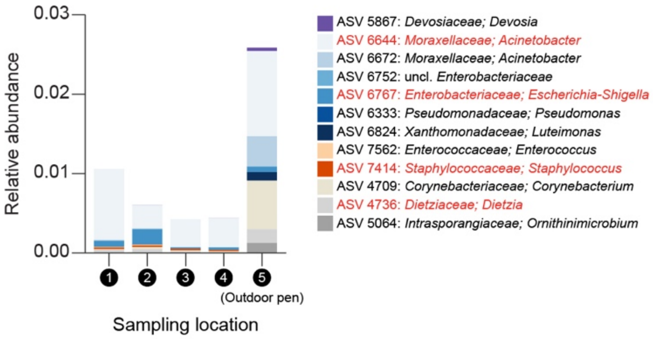


**Fig S16** Relative abundance of commonly shared ASVs in DCC-derived manure samples, by sampling location (*x*-axis; see Figure 1 for more information). Libraries derived from samples collected at a given sampling location were pooled across sampling dates for the bar graphs presented. Colored bars present the proportion of sequencing reads assigned to a given ASV. ASV taxonomy is shown at the genus level. Taxa highlighted in red were identified as commonly shared ASVs in both facilities.
